# Chromatin accessibility profiling in tissue sections by spatial ATAC

**DOI:** 10.1101/2022.07.27.500203

**Authors:** Enric Llorens-Bobadilla, Margherita Zamboni, Maja Marklund, Nayanika Bhalla, Xinsong Chen, Johan Hartman, Jonas Frisén, Patrik L Ståhl

## Abstract

Current methods for epigenomic profiling are limited in the ability to obtain genome wide information with spatial resolution. Here we introduce spatial ATAC, a method that integrates transposase-accessible chromatin profiling in tissue sections with barcoded solid-phase capture to perform spatially resolved epigenomics. We show that spatial ATAC enables the discovery of the regulatory programs underlying spatial gene expression during mouse organogenesis, lineage differentiation and in human pathological samples.

## Main text

In multicellular organisms, cells progressively acquire specialized gene expression programs according to their position within a tissue^1^. Cell type specific gene expression patterns result in part from the interaction between the transcriptional machinery and regulatory elements in the chromatin^2,3^, a process dysregulated in disease^4,5^. Multiple methods have been developed to integrate gene expression and chromatin accessibility measurements in single cells^6–8^. Single cell methods typically require tissue dissociation, and a wealth of spatial profiling methods have recently been developed to overcome this limitation, particularly on the transcriptome level^9^. However, we remain limited in our ability to interrogate chromatin accessibility with spatial resolution^10,11^.

We developed spatial ATAC to perform spatially resolved chromatin accessibility profiling in tissue sections. Spatial ATAC combines the assay for transposase-accessible chromatin and sequencing (ATAC-seq^12^) with tagmented DNA capture on a solid surface containing barcoded oligonucleotides, using an experimental platform analogous to our previous spatial transcriptomics approach^13^. First, we immobilize fresh frozen tissue sections onto barcoded slides and crosslink them to preserve chromatin structure during immunostaining. Immunostained sections are then imaged to register tissue coordinates and protein expression data. In the next step Tn5 transposition is performed directly in permeabilized sections to tagment open chromatin. With the help of a chimeric splint oligonucleotide, DNA tagments are hybridized to spatially barcoded surface oligonucleotides during gentle tissue digestion. Ligation to the splint and subsequent polymerase gap fill and extension generate open chromatin fragments carrying a spatial barcode and PCR handles that are used for generating a sequencing library (Fig. 1a).

**Figure 1.**
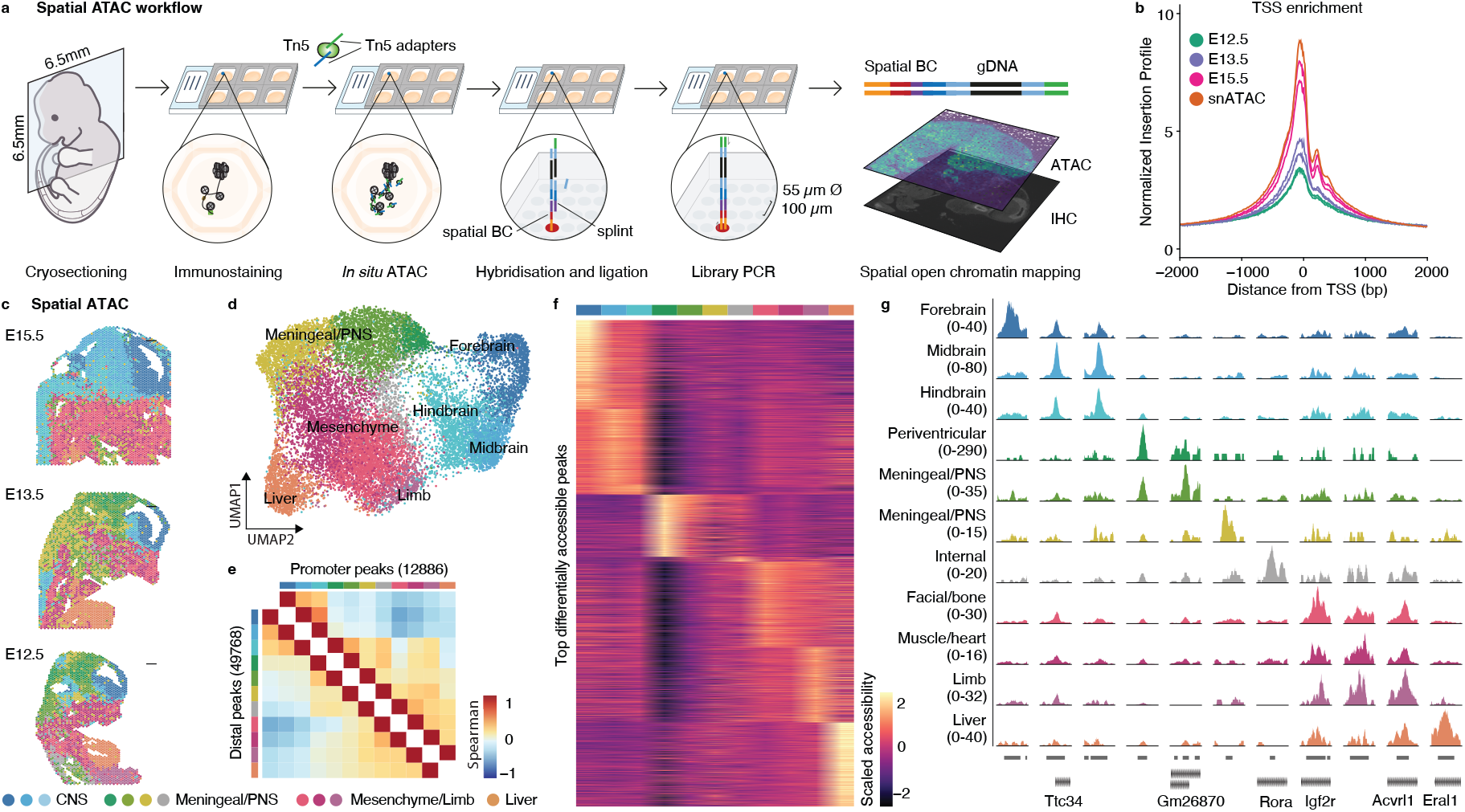
Workflow and spatial mapping of chromatin accessibility in mouse embryos. **a**. Schematic workflow of spatial ATAC. Transposition with Tn5 is performed on immunostained tissue cryosections immobilized on a barcoded slide. Transposed fragments are surface-captured using a splint oligonucleotide, which is ligated and extended to allow the generation of a spatially barcoded DNA library. **b**. Enrichment of ATAC-seq fragments around TSS in spatial ATAC performed on mouse embryos (E12.5, E13.5, E15.5) in comparison with single-nucleus ATAC-seq from 10X genomics (E18; snATAC). **c**. Clustering of spatial ATAC open chromatin fragments projected on their spatial location. **d**. UMAP of all spots from mouse embryo sections colored by cluster as in c. **e**. Cluster-wise correlation of the accessibility of the top 25% variable promoter (+1000, -100bp from TSS) and distal peaks. **f**. Heatmap showing scaled accessibility of the top differentially accessible peaks per cluster. **g**. Genome tracks showing normalized spatial ATAC-seq fragment density for peaks showing cluster-specific accessibility. Cluster colors are consistent from c-g. Scale bars are 500 µm.

We performed spatial ATAC on replicate tissue sections from three stages of mouse gestational development (embryonic days E12.5, E13.5 and E15.5). Spatially barcoded open chromatin fragments showed high enrichment around transcriptional start sites (TSS) as well as nucleosome periodicity, hallmarks of ATAC-seq (Fig. 1b and Extended Data Fig. 1). We captured a median of 6100, 3100 and 7100 unique fragments per 55 µm spot, with 14, 15 and 18% overlapping TSS in E12.5, E13.5 and E15.5 sections respectively. These metrics are within the range of reference single-nucleus ATAC-seq data from E18 mouse brain (Extended Data Fig. 1a-d). Additionally, the aggregate distribution of fragments across the genome showed a very high concordance with reference bulk datasets from ENCODE^14^ (Extended Data Fig. 1e). We next created a peak spatial barcode count matrix using a common reference peak set across sections that were analyzed by latent semantic indexing (LSI) and uniform manifold approximation and projection (UMAP) for dimensionality reduction^15^. Unsupervised clustering identified 11 main clusters, which when projected in their original spatial coordinates revealed a high concordance with anatomical landmarks and were consistent not only across replicate sections but also across developmental stages (Fig. 1c-d and Extended Data Fig. 2). This clustering agreed with spatial-aware non-negative matrix factorization (NMF) dimensionality reduction and clustering^16^, suggesting that spatial location is a major source of variation in chromatin accessibility across and within developing tissues (Extended Data Fig. 3a-d). As expected, the dataset structure reflected variation in the accessibility of promoters and a larger set of distal peaks (Fig. 1e). Using differential accessibility analyses we found 18,000 differentially accessible peaks that showed specific patterns of accessibility across developing tissues (Fig. 1f-g).

**Figure 2.**
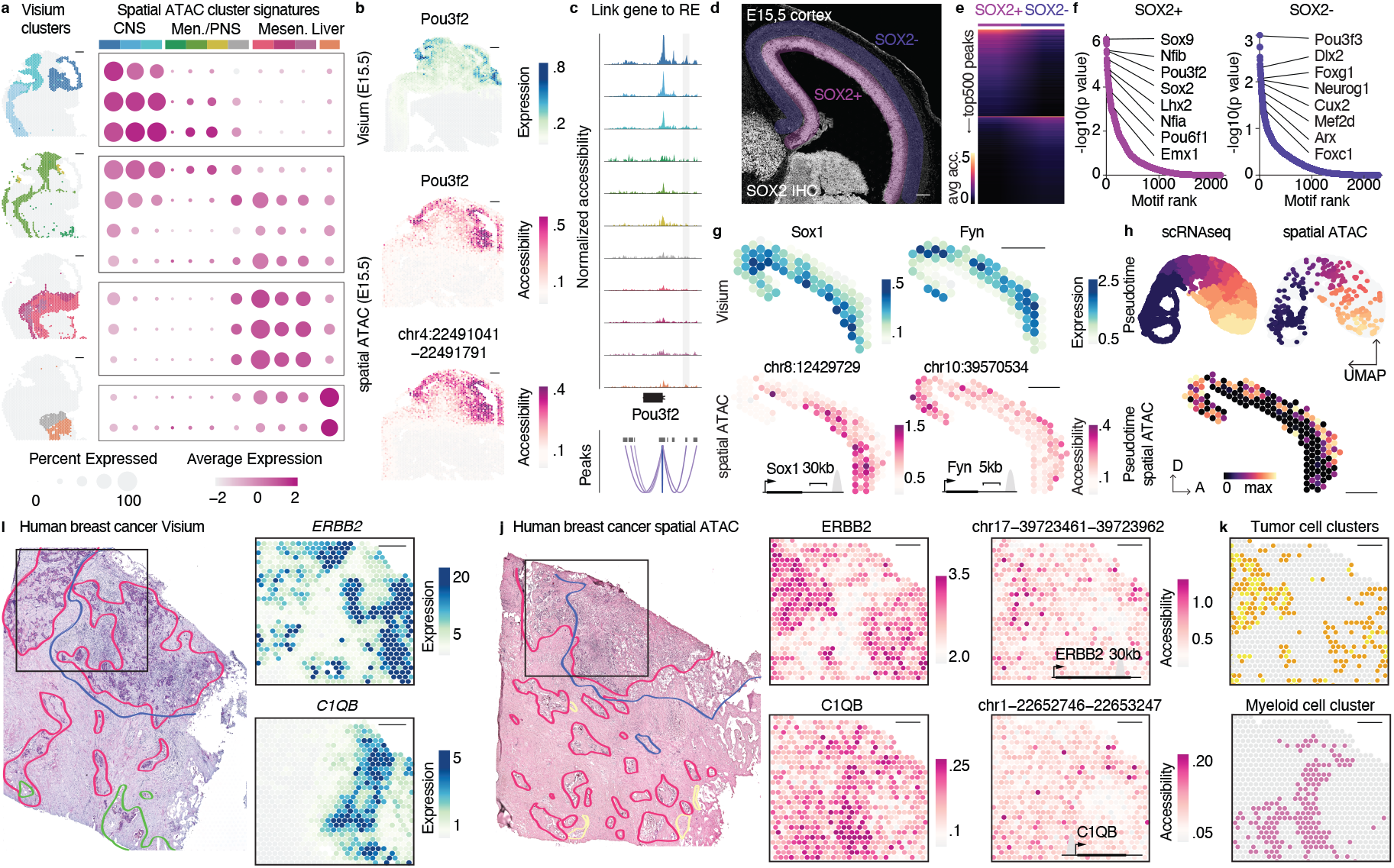
Spatial ATAC uncovers spatiotemporal patterns of regulatory element accessibility underlying gene expression. **a**. Visium gene expression signature scores for differentially accessible genes in spatial ATAC clusters. Visium clusters are shown on the left on an E12.5 section for reference. **b**. Concordance between Pou3f2 expression (top, cyan) and gene activity and accessibility of a co-accessible distal regulatory element (magenta). **c**. Genomic track and co-accessibility scores for peaks near the locus of the CNS marker Pou3f2. The distal element shown in b is highlighted in gray and tracks are color-coded according to spatial ATAC clusters. **d**. Cortex inset of a SOX2-immunostained E15.5 spatial ATAC sagittal section. Selected SOX2+ (progenitor) and SOX2- (neuronal) regions are highlighted. **e**. Top 500 differentially accessible peaks by fold change in SOX2+ and SOX2- regions of the developing cortex. **f**. Motif enrichment analysis performed on the top 500 peaks by region. Selected top motifs for transcription factors expressed in the region are highlighted. **g**. Accessibility (spatial ATAC; magenta) and expression of the nearest gene (Visium; cyan) for loci enriched in progenitor (Sox1) or neuronal (Fyn) regions. **h**. UMAP of integrated single-cell RNA-seq and spatial ATAC from the E15.5 developing cortex colored by pseudotime and split by technology. At the bottom, pseudotime scores are projected onto their spatial locations in a spatial ATAC E15.5 section. **i**. HE image of a breast cancer section processed using Visium with overlaid pathologist annotations. On the right, expression of ERBB2 (HER2) and myeloid cell marker C1QB in the boxed inset. **j**. Annotated HE image of an adjacent (200 µm) section processed using spatial ATAC. On the right, accessibility of the ERBB2 locus, C1QB locus and two associated regulatory regions in the boxed inset. **k**. Spatial interaction between tumor cell and myeloid cell clusters at the tumor interface. Pathology: red, invasive cancer; blue, tumor infiltrating lymphocytes; green, intravascular cancer; yellow, normal gland. Scale bars are 500 µm.

We next computed gene activities (i.e., accessibility at gene locus and promoter), which revealed 2000 differentially accessible genes between clusters that were enriched for gene ontology terms characteristic of the respective tissue region (Extended Data Fig. 4). For example, central nervous system (CNS) clusters showed increased accessibility in genes known to be involved in neurogenesis (e.g., Sox1, Foxg1, Notch1). Bone and muscle mesenchyme clusters showed increased accessibility in myofiber, collagen, and TGF-b signaling genes (e.g., Myh9, Col1a1, Smad3) while the fetal liver cluster was characterized by the accessibility of genes involved in hematopoiesis (e.g., Hba-a1, Tal1, Sptb).

Next, we sought to integrate spatial ATAC with Visium spatial transcriptomics. We performed Visium on tissue sections from the same developmental stages, which showed regionally consistent clustering (Extended Data Fig. 5) and genes found as differentially accessible using spatial ATAC showed higher expression in the corresponding Visium cluster (Fig. 2a). Unsupervised denoising and imputation methods have been developed to account for the intrinsic sparsity of single-cell transcriptomics and ATAC-seq data which improve visualization and feature-to-feature correlation^17,18^. We applied a denoising deep count autoencoder to our spatial ATAC and Visium datasets^18^, which increased signal to noise in feature visualization while preserving clustering structure (Extended Data Fig. 5). To identify putative regulatory elements underlying spatial patterns of gene expression, we performed peak co-accessibility analyses which identified 6000 peaks linked to cluster marker genes. With this strategy, we identified individual distal regulatory elements whose accessibility correlated to gene expression across tissues (Extended Data Fig. 6) and agreed with enhancer reporter assays (Extended Data Fig. 7). To gain further insight into regulatory programs underlying gene expression, we performed motif enrichment analysis on these cluster-specific distal peaks. We found that the most enriched motifs in CNS clusters corresponded to well characterized proneural transcription factors (e.g., Neurog1, Neurod1, Ascl1). Conversely, motifs enriched in mesenchymal regulatory elements corresponded to factors known to be involved in bone and muscle development (e.g., Smad3, Twist1, Myog), while liver-specific distal regulatory elements were highly enriched in binding sites for Tal1 and Gata transcription factors, consistent with their role in hematopoiesis (Extended Data Fig. 6d).

To evaluate whether spatial ATAC could identify regulatory programs underlying lineage differentiation within a developing tissue, we focused on the cerebral cortex at E15.5, a well characterized structure in which SOX2+ progenitors in the subventricular zone generate neurons that migrate to upper cortical layers^19^. Based on SOX2 immunostaining, we selected progenitor- and neuron-rich spots and performed motif enrichment on the top differentially accessible peaks (Fig. 2d-f). We identified cortical progenitor (e.g., Sox2, Lhx2, Emx1) and neuronal (e.g., Neurog1, Cux2) transcription factors among the top enriched motifs in the respective clusters (Fig. 2f). Further, we could link regulatory elements to the nearest genes that showed the corresponding patterns of layer-specific gene expression (Fig. 2g). Next, we integrated the cortex spatial ATAC spots with single cell RNA-seq data from the same developmental stage^20^. Using the integrated dataset, we calculated pseudotime scores along the neuronal differentiation trajectory, which aligned single cells and spatial ATAC spots and recapitulated the inside-out differentiation trajectory of the developing cortex (Fig. 2h).

Finally, we applied spatial ATAC to human breast cancer, a tumor type of widespread public health concern in which pathological classification informs therapy decisions^21^. We profiled adjacent sections using Visium and spatial ATAC. Spatial ATAC clustering and marker expression aligned with pathologist annotations, agreed with Visium clustering, and could readily identify HER2-positive regions, their associated non-coding region accessibility, and the presence of myeloid cells in the immediate tumor microenvironment (Fig. 2i-k, Extended Data Fig. 8-10).

Our spatial ATAC platform is readily implementable through common laboratory workflows and offers the possibility for integration with other ‘omics modalities. We envision that spatial ATAC will enable spatial non-coding functional genomics, while being instrumental in the identification of regulatory elements for specific cell targeting in gene therapy and the study of gene regulatory networks in development and disease.

## Acknowledgements

A. Andersson for initial help and advice with the analyses. L. Larsson, M. Lukoseviciute and C. Engblom for helpful discussions. V. Kumar for help adapting Cell Ranger. P. Backhaus for help in embryo harvesting. G. Winberg, H. Lönnqvist, M. Hagemann-Jensen and R. Sandberg for help preparing Tn5 and access to NextSeq sequencing. We thank National Genomics Infrastructure (NGI), Sweden for providing infrastructure support. The data were analyzed using resources provided by SNIC through the Uppsala Multidisciplinary Center for Advanced Computational Science (SNIC/UPPMAX).

This work was supported by grants from the Swedish Research Council, the Strategic Research Programme in Stem Cells and Regenerative Medicine at Karolinska Institutet (StratRegen), and the Torsten Söderbergs Stiftelse.

## Author contributions

E.L.B., J.F. and P.L.S. conceived the project. E.L.B, M.M. and N.B. performed the experiments. M.Z., E.L.B. conducted the analyses and visualizations. X.C and J.H. provided cancer samples and pathology annotations. E.L.B. wrote the manuscript with input from all the authors. J.F. and P.L.S. acquired funding and supervised the project.

## Competing interests

E.L.B., M.Z., M.M., N.B., J.F. and P.L.S. are scientific consultants to 10x Genomics, which holds IP rights to the spatial technology.

## References

1. Nitzan, M., Karaiskos, N., Friedman, N. & Rajewsky, N. Gene expression cartography. Nature 576, 132–137 (2019).

2. Klemm, S. L., Shipony, Z. & Greenleaf, W. J. Chromatin accessibility and the regulatory epigenome. Nat Rev Genet 20, 207–220 (2019).

3. Shen, Y. et al. A map of the cis-regulatory sequences in the mouse genome. Nature 488, 116–120 (2012).

4. Corces, M. R. et al. The chromatin accessibility landscape of primary human cancers. Science 362, eaav1898 (2018).

5. Ge, Y. et al. Stem Cell Lineage Infidelity Drives Wound Repair and Cancer. Cell 169, 636-650.e14 (2017).

6. Satpathy, A. T. et al. Massively parallel single-cell chromatin landscapes of human immune cell development and intratumoral T cell exhaustion. Nature biotechnology 37, 925–936 (2019).

7. Ma, S. et al. Chromatin Potential Identified by Shared Single-Cell Profiling of RNA and Chromatin. Cell 183, 1103-1116.e20 (2020).

8. Chen, S., Lake, B. B. & Zhang, K. High-throughput sequencing of the transcriptome and chromatin accessibility in the same cell. Nat Biotechnol 37, 1452–1457 (2019).

9. Palla, G., Fischer, D. S., Regev, A. & Theis, F. J. Spatial components of molecular tissue biology. Nat Biotechnol 40, 308–318 (2022).

10. Deng, Y. et al. Spatial-CUT&Tag: Spatially resolved chromatin modification profiling at the cellular level. Science 375, 681–686 (2022).

11. Thornton, C. A. et al. Spatially mapped single-cell chromatin accessibility. Nat Commun 12, 1274 (2021).

12. Buenrostro, J. D., Giresi, P. G., Zaba, L. C., Chang, H. Y. & Greenleaf, W. J. Transposition of native chromatin for fast and sensitive epigenomic profiling of open chromatin, DNA-binding proteins and nucleosome position. Nature methods 10, 1213–1218 (2013).

13. Ståhl, P. L. et al. Visualization and analysis of gene expression in tissue sections by spatial transcriptomics. Science (New York, NY) 353, 78–82 (2016).

14. Gorkin, D. U. et al. An atlas of dynamic chromatin landscapes in mouse fetal development. Nature 583, 744–751 (2020).

15. Stuart, T., Srivastava, A., Madad, S., Lareau, C. A. & Satija, R. Single-cell chromatin state analysis with Signac. Nat Methods 18, 1333–1341 (2021).

16. Bergenstråhle, J., Larsson, L. & Lundeberg, J. Seamless integration of image and molecular analysis for spatial transcriptomics workflows. BMC Genomics 21, 482 (2020).

17. Li, Z. et al. Chromatin-accessibility estimation from single-cell ATAC-seq data with scOpen. Nat Commun 12, 6386 (2021).

18. Eraslan, G., Simon, L. M., Mircea, M., Mueller, N. S. & Theis, F. J. Single-cell RNA-seq denoising using a deep count autoencoder. Nat Commun 10, 390 (2019).

19. Greig, L. C., Woodworth, M. B., Galazo, M. J., Padmanabhan, H. & Macklis, J. D. Molecular logic of neocortical projection neuron specification, development and diversity. Nature Reviews Neuroscience 14, 755–769 (2013).

20. La Manno, G. et al. Molecular architecture of the developing mouse brain. Nature 596, 92–96 (2021).

21. Wu, S. Z. et al. A single-cell and spatially resolved atlas of human breast cancers. Nat Genet 53, 1334–1347 (2021).

