## Supplemental material for "Chromatin accessibility profiling in tissue sections by spatial ATAC"

**Extended Data Figure 1**

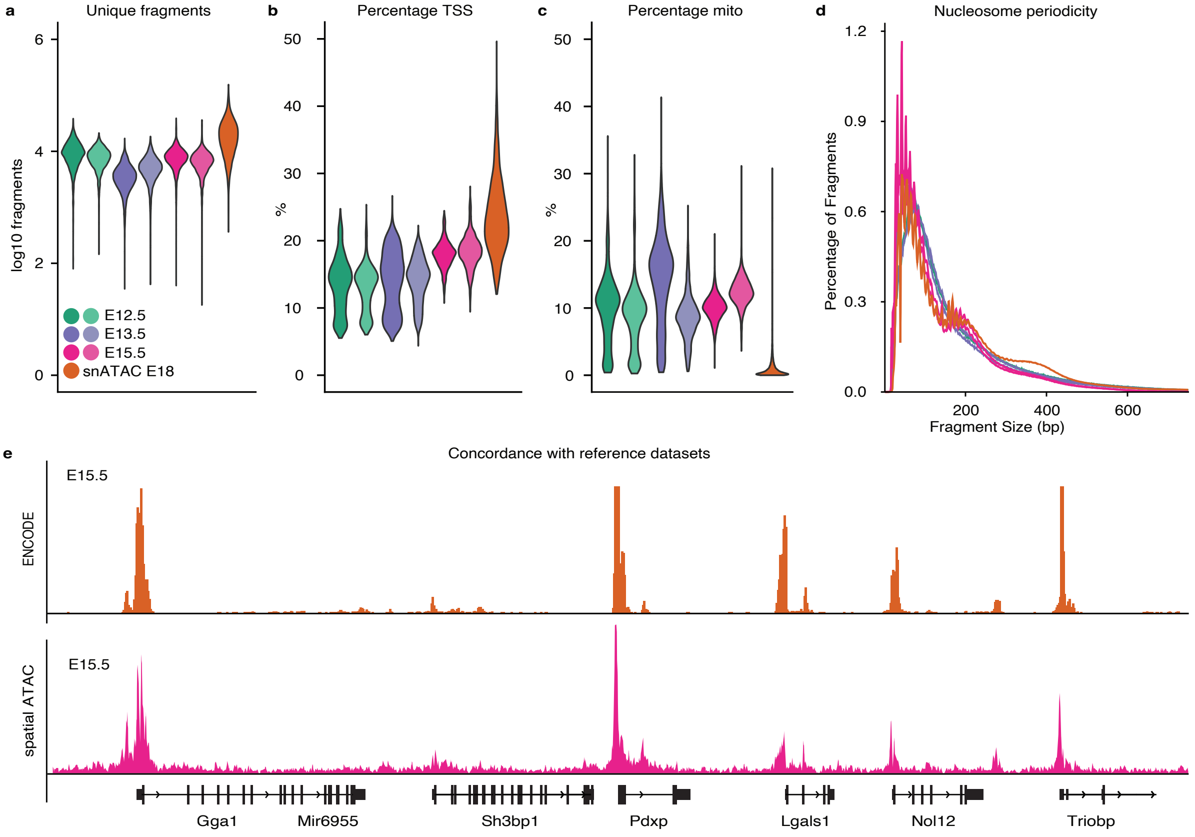

**Extended Data Figure 1**. Quality control metrics of spatial ATAC. **a-c.** Violin plot showing unique fragments per spot, percentage TSS fragments and percentage mitochondrial reads in embryo sections processed using spatial ATAC and single-nucleus ATAC-seq from 10X Genomics (i.e., a flash frozen cortex, hippocampus, and ventricular zone from embryonic mouse brain; E18). **d**. Fragment size histogram colored by technology and replicate as in a. **e.** Genome ATAC-seq coverage tracks from bulk ENCODE E15.5 ATAC-seq datasets (brain and facial prominence) and aggregate signal from two spatial ATAC E15.5 sections.

Extended Data Figure 2

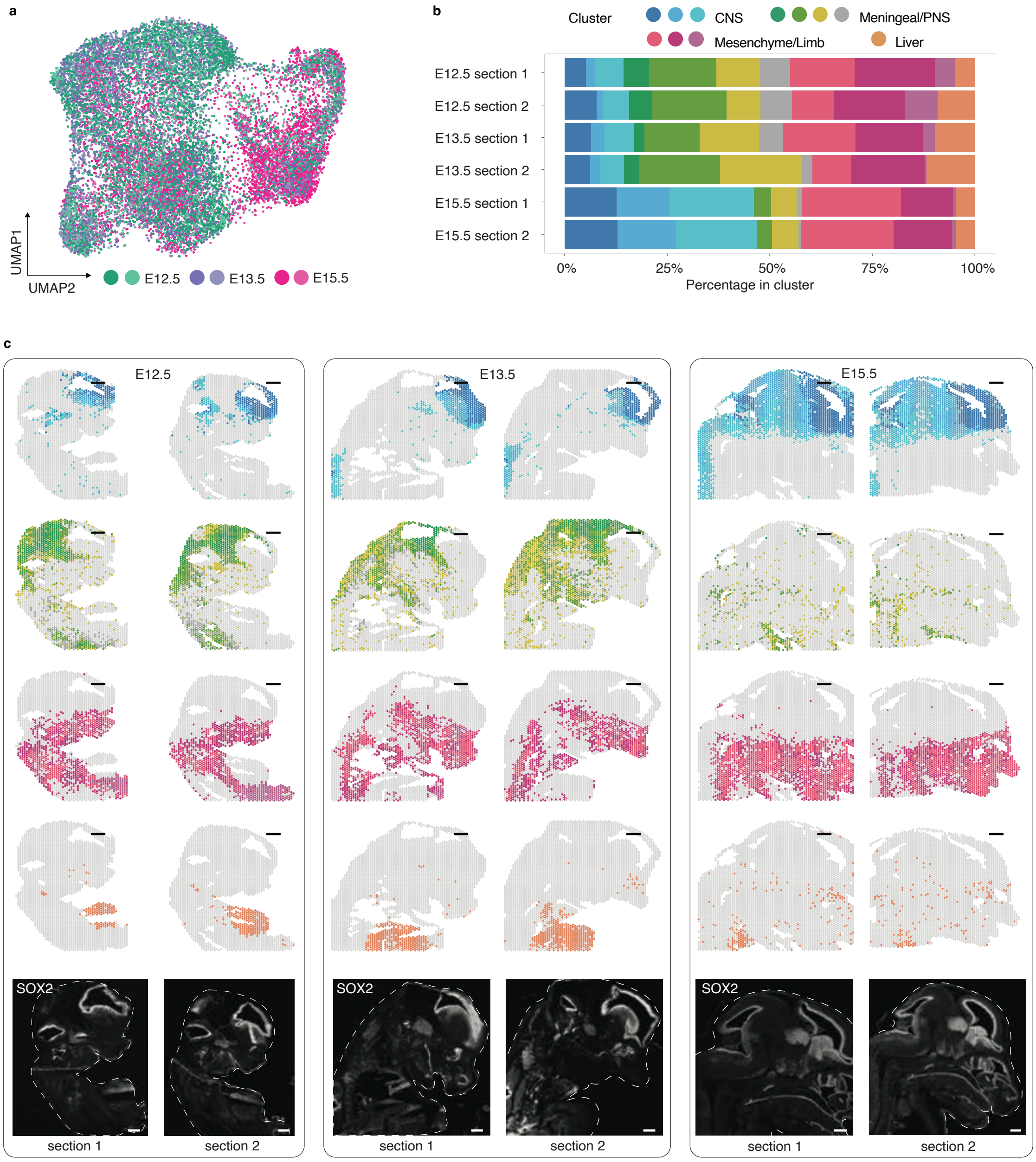

**Extended Data Figure 2**. spatial ATAC in mouse embryos**. a.** UMAP embedding corresponding to Fig. 1d but colored by embryonic age and section replicate. **b.** Cluster proportions across embryo sections. **c.** Cluster families as in Fig. 1a for all sections analyzed. SOX2 immunostaining for the respective section at the bottom. Scale bars are 500µm.

**Extended Data Figure 3**

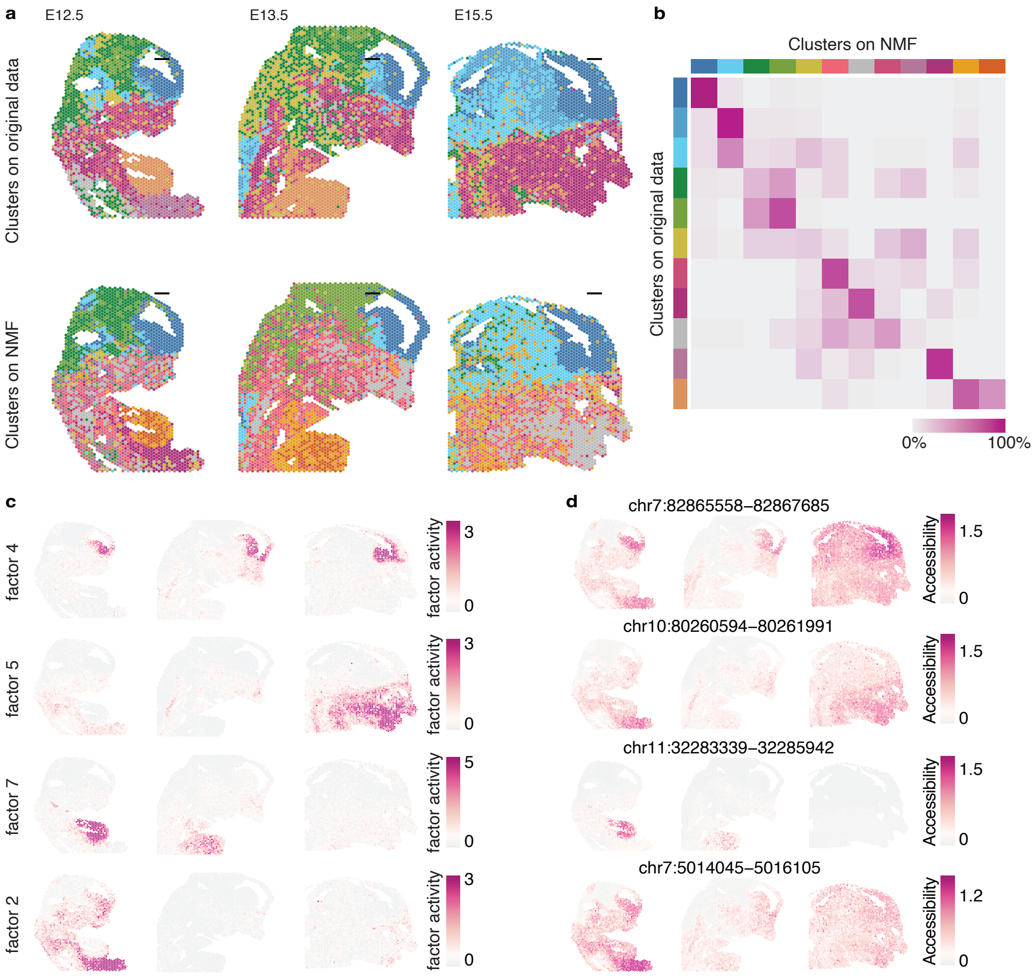

**Extended Data Figure 3.** Clustering of spatial ATAC data with spatially aware factor analysis. **a.** Spatial ATAC clusters based on LSI (top) or NMF (bottom) for dimensionality reduction. **b**. Heatmap displaying the percentage of spots assigned to LSI- or NMF-computed clusters. **c.** Spatial activity plots for selected factors enriched in forebrain, facial prominence, liver, and limb. **d.** Examples of the most contributing peaks for each factor represented in c. Scale bars are 500 μm.

**Extended Data Figure 4**

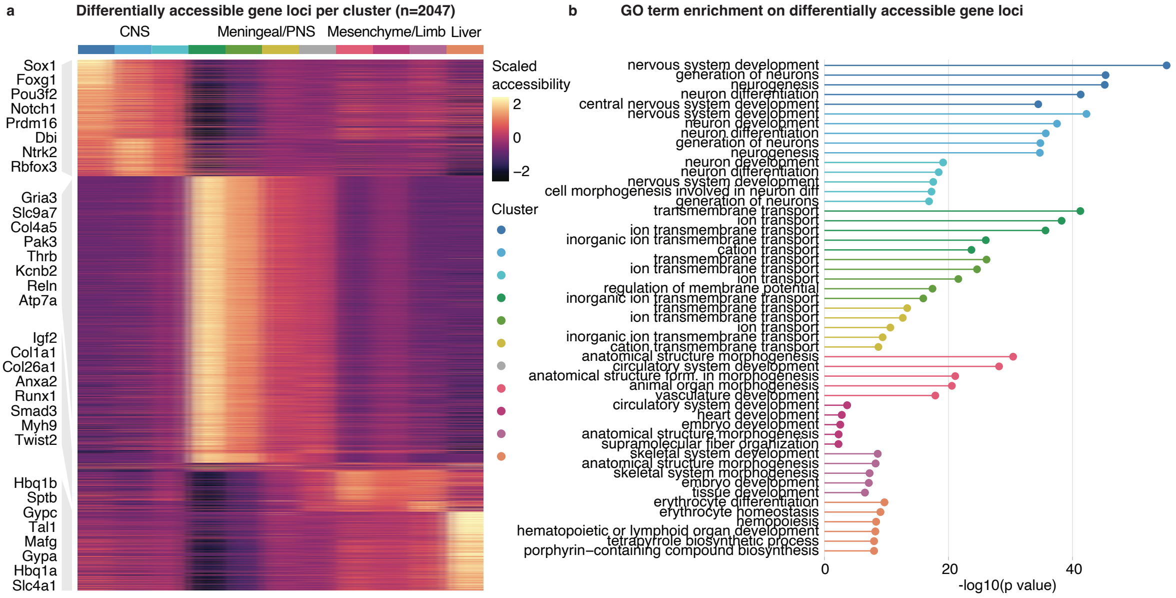

**Extended Data Figure 4.** Cluster marker genes and gene ontology analysis. **a.** Heatmap showing scaled accessibility for the top differentially accessible genes (gene body + promoter) across clusters. Relevant markers are highlighted. **b.** Gene ontology enrichment analysis of the top marker genes colored by cluster.

Extended Data Figure 5

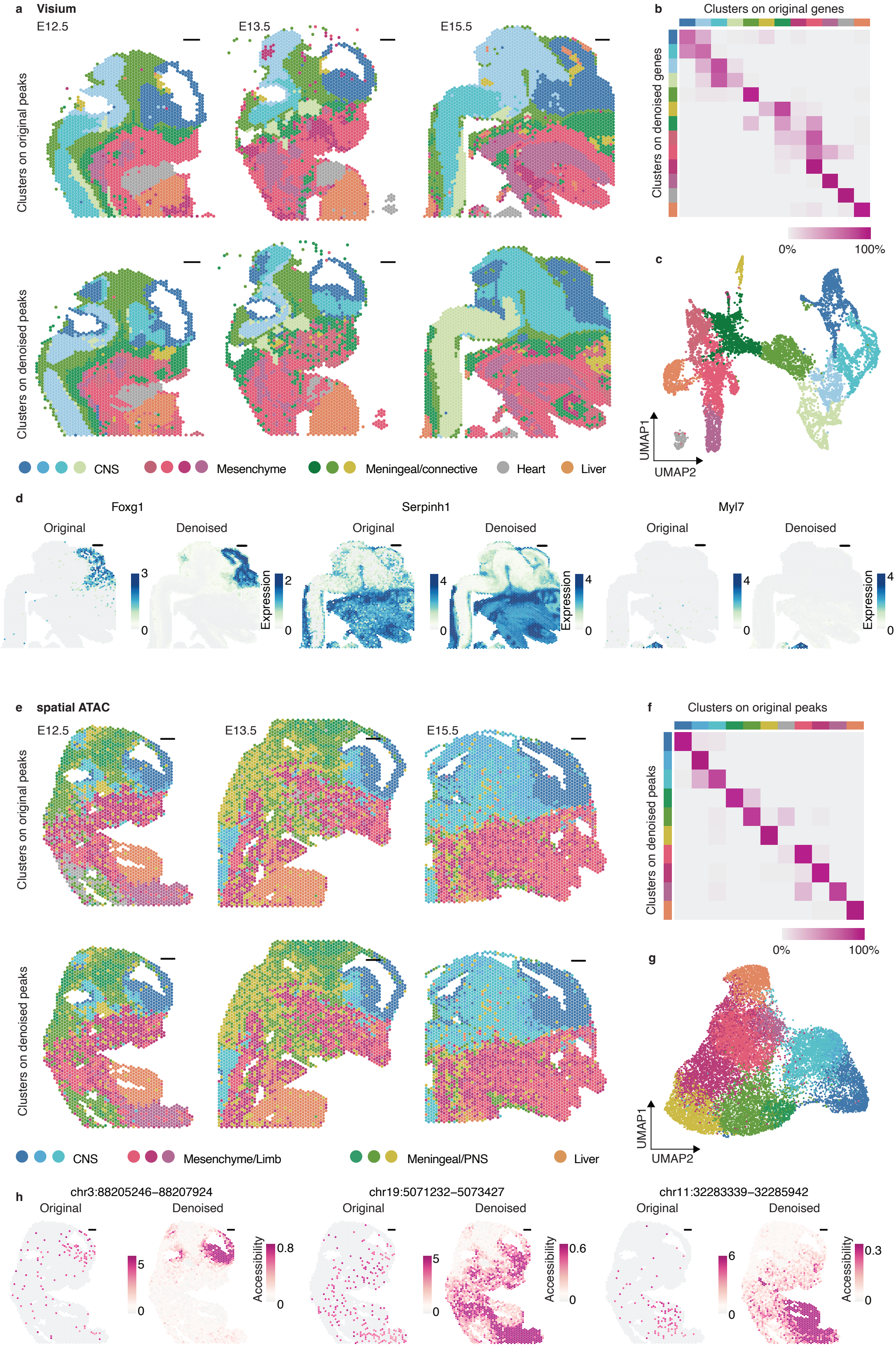

**Extended Data Figure 5.** Deep count autoencoder denoising of Visium and spatial ATAC **a**. Visium clusters based on original or denoised gene counts. **b**. Heatmap displaying the percentage of spots being assigned to the clusters obtained from original or denoised Visium data. **c**. UMAP on denoised peaks colored by cluster. **d**. Visualization of gene expression normalized counts before and after denoising on E15.5 sections. **e**. Clusters based on original or denoised spatial ATAC peak counts. **f**. Heatmap displaying the percentage of spots being assigned to the clusters obtained from original or denoised spatial ATAC data **g**. UMAP on denoised peaks colored by cluster. **h**. Visualization of normalized peak accessibility before and after denoising on E12.5 sections. Scale bars are 500 μm.

Extended Data Figure 6

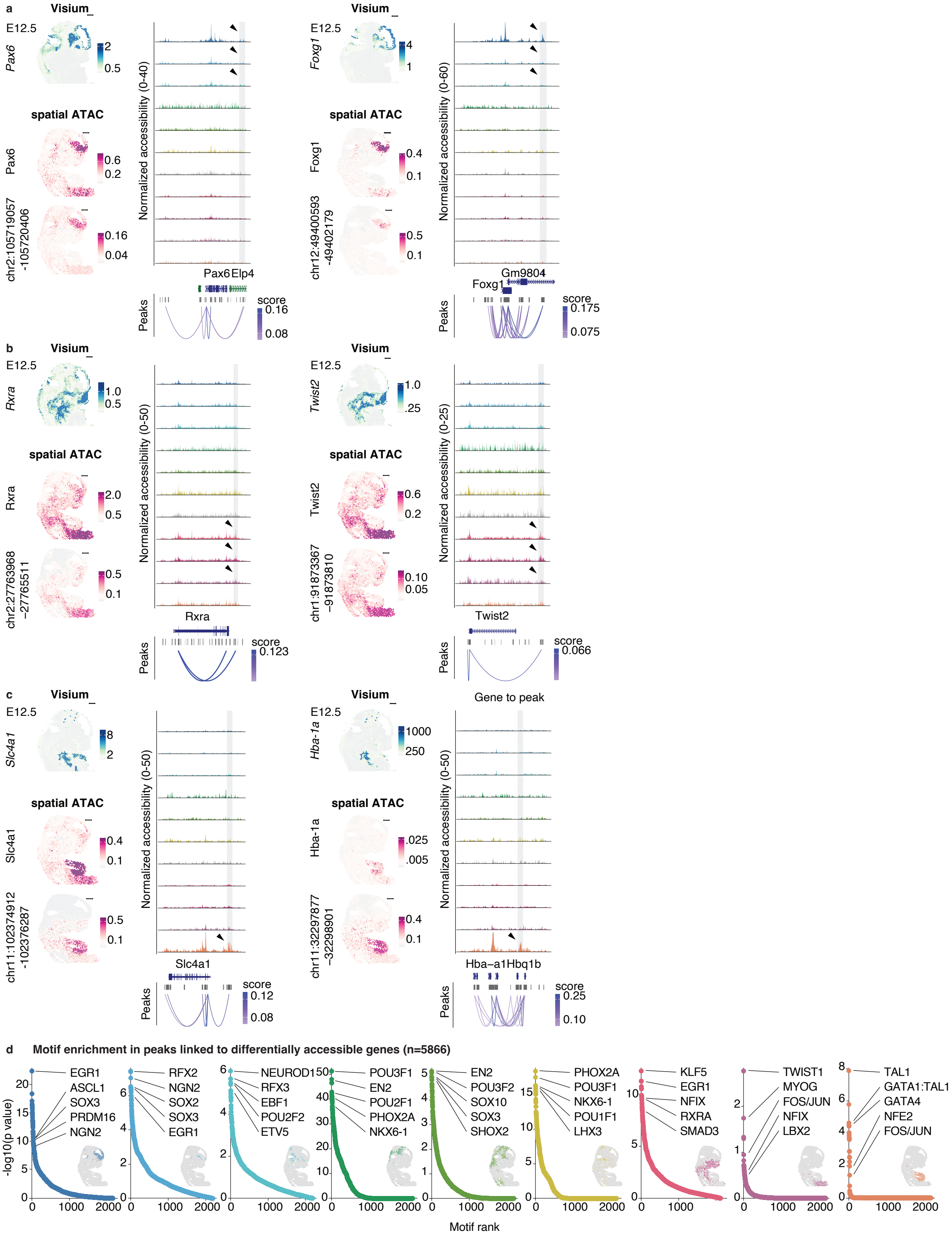

**Extended Data Figure 6.** Gene regulatory programs during mouse organogenesis. **a**. Visium expression, spatial ATAC gene activity and regulatory element accessibility at E12.5 for CNS/Forebrain markers Pax6 and Foxg1. The respective linked regulatory element is shown in gray. **b.** Visium expression, spatial ATAC gene activity and regulatory element accessibility for Mesenchyme/Limb markers Rxra and Twist2. The respective linked regulatory element is shown in gray. **c.** Visium expression, spatial ATAC gene activity and regulatory element accessibility for liver markers Slc4a1 and Hba-a1. The respective linked regulatory element is shown in gray. Arrowheads point to clusters for which the regulatory element is most accessible. **d.** Motif enrichment rank plots for cluster-specific distal elements. Selected top non-redundant transcription factor motifs are highlighted. Scale bars are 500 μm.

Extended Data Figure 7

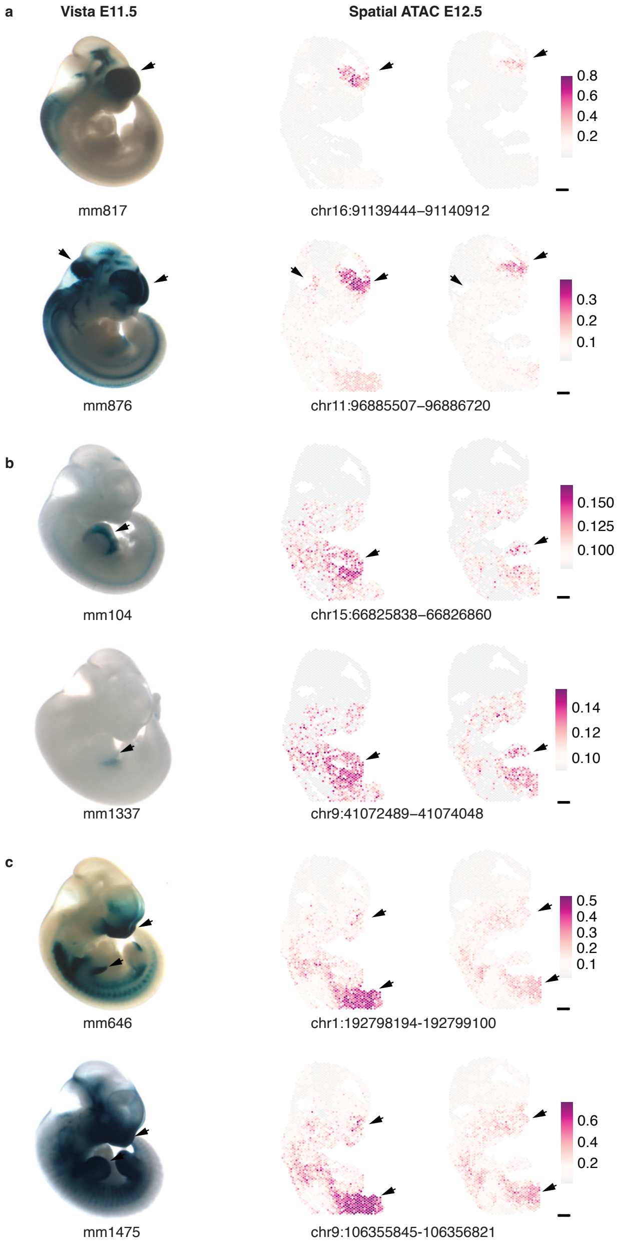

**Extended Data Figure 7.** Validation of spatial ATAC regulatory element accessibility. **a**. Vista enhancer reporter expression for two CNS elements overlapping with differentially accessible spatial ATAC peaks. **b**. Vista enhancer reporter expression for two liver elements overlapping with differentially accessible spatial ATAC peaks. **c**. Vista enhancer reporter expression for two limb elements overlapping with differentially accessible spatial ATAC peaks. Reporter images were obtained from https://enhancer.lbl.gov/ Scale bars are 500 µm.

**Extended Data Figure 8**

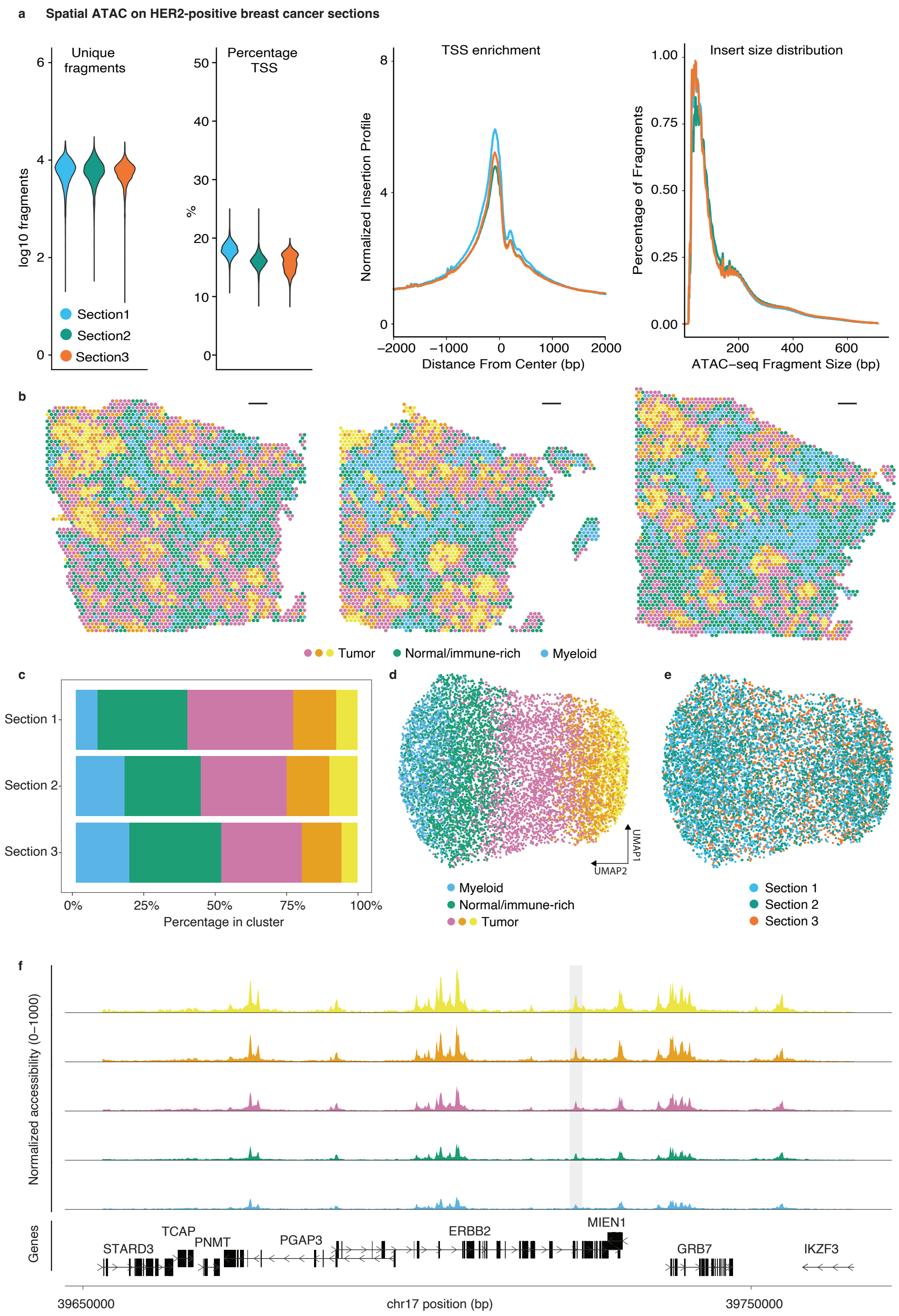

**Extended Data Figure 8.** Spatial ATAC on human HER2-positive breast cancer. **a.** On the left, violin plot showing unique fragments per spot and percentage TSS fragments in three adjacent sections processed using spatial ATAC. On the right, TSS enrichment and insert size distribution. **b**. Spatial ATAC clustering reveals tumor, immune-rich and normal tissue regions. **c.** Cluster percentage across sections. **d-e**. UMAP embedding on spatial ATAC peaks color-coded by cluster or tissue section. Fragment size histogram color-coded by technology and replicate as in a. **f.** Genome tracks showing normalized spatial ATAC-seq fragment density around the HER2 (ERBB2) locus colored by cluster. The gray area marks a previously described gene body enhancer (Liu et al. 2018) shown in Fig. 2j. Scale bars are 500 μm.

**Extended Data Figure 9**

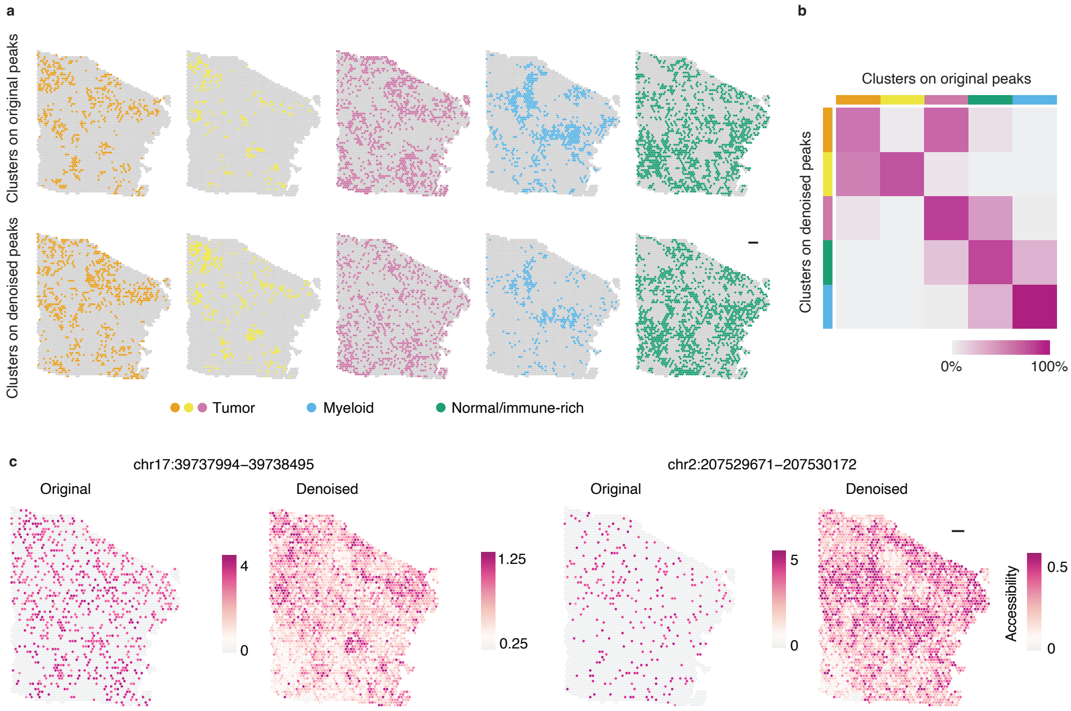

**Extended Data Figure 9.** Deep count autoencoder denoising of spatial ATAC breast cancer data **a**. Clusters based on original or denoised peak counts. **b**. Heatmap displaying the percentage of spots being assigned to the clusters obtained from original or denoised spatial ATAC data. **c**. Visualization of peak accessibility scores before and after denoising on one example section. Scale bars are 500 μm.

Extended Data Figure 10

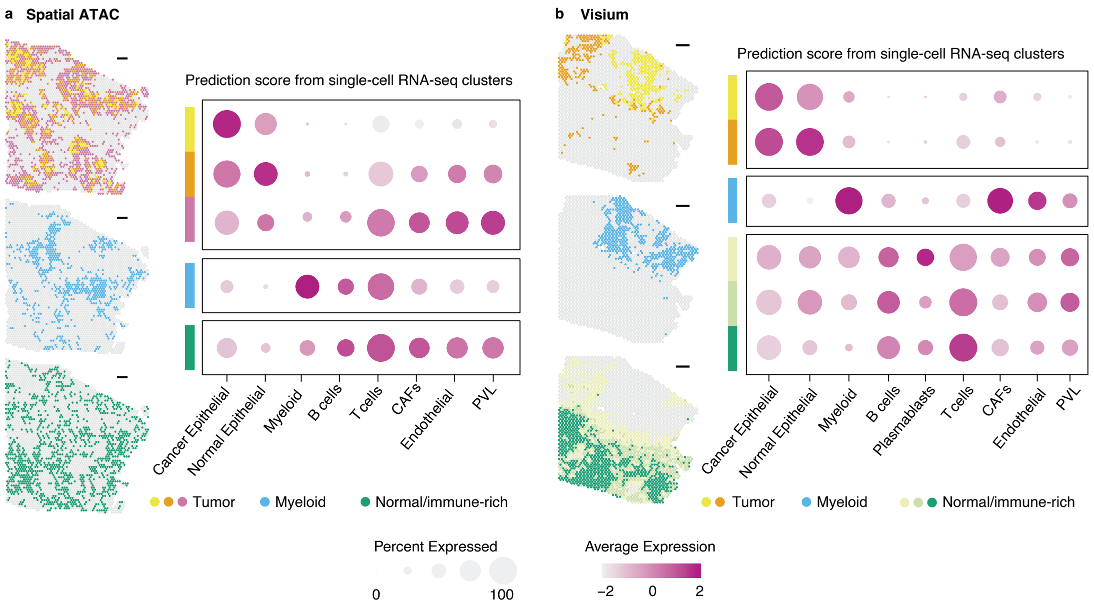

**Extended Data Figure 10.** Multimodal integration of single-cell RNA-seq with spatial ATAC and Visium on breast cancer sections. **a.** Prediction scores from scRNA-seq signatures in spatial ATAC clusters reveal cell composition differences across clusters. **b**. Same as in a but for Visium clusters. Scale bars are 500 μm.

### Material and Methods

#### Animal tissue processing

Time pregnant C57BL/6 mice were purchased from Janvier and were sacrificed by cervical dislocation at embryonic day 12.5, 13.5 or 15.5 for embryo harvesting. All experimental procedures were carried out in accordance to the Swedish and European Union guidelines and approved by the institutional ethical committee (Stockholms Norra Djurförsöksetiska Nämnd) under ethical permit numbers N155/16 and 20785/2020.

The tissues were harvested on ice-cold PBS and snap frozen in OCT (Tissue-Tek, 4583) blocks in a dry ice-isopentane bath at -60°C and stored at -80°C until sectioned.

#### Collection of breast cancer patient tumor samples

Breast cancer tissues were obtained from the Department of Clinical Pathology and Cancer Diagnostics at Karolinska University Hospital, Stockholm, Sweden. Experimental procedures and protocols were approved by the regional ethics review board (Etikprövningsnämnden) in Stockholm (2016/957-31, amendment 2017/742-32 and 2021-00795), and informed consent was obtained from the participating patient.

The samples were obtained from a breast tumor removed from a patient with treatment-naive invasive ductal carcinoma. The tumor was divided into several regions and collected freshly by a pathologist depending on the size of the tumor. From each region, tissue was isolated for direct embedding in OCT, followed by immediate freezing and storage at -80°C until further analysis. Histological evaluations of the patient tumor were performed by pathologists for diagnostic purposes: tumor characteristics, including grade, size, hormone receptor, HER2, and Ki67 status are presented in Table S3.

#### Spatial ATAC

Cryosections were cut on a cryostat (Leica, NX70) at a 10 μm thickness and placed on spatially barcoded OMNI glass slides (10X Genomics). In brief, each OMNI array slide contained eight capture areas, each covered by 5000 barcoded spots with a diameter of 55 μm and 100 μm between spots. Each spot contained millions of DNA oligonucleotides encoding a 16nt spatial barcode, serving as x and y coordinate, a PCR handle for library amplification, a 12nt UMI, and a 7nt generic capture sequence used for splint oligonucleotide hybridisation (see Table S1).

Slides were first heated at 37°C for 1 min to adhere the tissue to the slide. Then, the sections were crosslinked in freshly prepared methanol-free 0.5% formaldehyde (Polysciences, 18814) diluted in DPBS for 10 minutes at room temperature, followed by rinsing in 500 mM Tris-HCl pH8 (Thermo, AM9856) to quench formaldehyde. After dipping the slide in DPBS three times, the sections were immunostained as follows: The tissue sections were blocked by incubation for 5 minutes with staining buffer (DPBS containing 5% Donkey serum, 0.1% NP-40 – Thermo 28324- and 0.005% Digitonin – Promega G9441). The staining buffer was then removed, and the primary antibody dilution added (antibodies: rabbit anti-SOX2 Merck 5603; goat anti-SOX9 R&D 3075; anti-nuclear antigen Novus 235-1) and incubated at room temperature for 30 minutes. Then, washing was performed 2 times with staining buffer for 3 minutes each, followed by addition of Alexa 647-conjugated secondary antibody dilution (Thermo 31573 or 21447), and incubation at room temperature for 15 minutes. Then, washing was performed 3 times with staining buffer for 3 minutes each, and finally pipette washed with DPBS once. The slides were then spin-dried, covered with 85% glycerol, mounted with a coverslip, and imaged in a Zeiss LSM 700 (x10 magnification) confocal or in a Metafer VSlide system (x20 magnification) epifluorescence microscope to record tissue coordinates and capture area fiducials. The images were processed with the VSlide software (v1.0.0) or with Fiji [1].

After image acquisition, the glycerol was removed by dipping in DPBS and a layer of isopropanol was then added to the arrays, decanted, and air-dried. The slide was then re-

hydrated in DPBS followed by ATAC permeabilization (0.01% digitonin, 0.1% Tween-20, 0.1% NP-40, 10 mM Tris-HCl pH7.4, 10 mM NaCl, 3 mM MgCl2) at room temperature for 10 minutes.
Custom Tn5 transposomes (30 µM) were assembled using Nextera adapter oligonucleotides A and B (see Table S1) according to [2]. Tagmentation was performed according to OMNI ATAC-seq [3] at 37°C for 1 h under gentle shaking (300 rpm every 5 minutes) using 2 µl Tn5 in tagmentation mix (25 µl 2x TD buffer, 16,5 µl DPBS, 0.5 µl 1% digitonin, 0.5 µl 10% Tween-20). To stop the tagmentation and strip the transposase from DNA, sections were incubated with 50 mM EDTA while ramping down to 30°C for 10 minutes. To hybridise the tagments to the barcoded surface oligonucleotides, we then incubated the sections with a 2 µM solution of splint oligonucleotide (in 3X SSC buffer containing 0.01% Triton-X100, 0.8 µg/µl Proteinase K and 2.5% PEG8000) overnight at 30°C. Next, the sections were rinsed in 2X NEB 2.1 buffer and subsequently incubated with ligation and polymerization solution (1X NEB2.1 containing 3U T4 DNA polymerase, 2000U T4 DNA ligase, 100 uM dNTPs, 1 mM ATP, all from NEB) and incubated at 18°C for 4h. Tissue removal was then performed using 2 mg/ml Proteinase K in PKD-buffer (Qiagen), and incubated at 56°C for 30 minutes (shaking at 300 rpm). The slides were then sequentially washed in 2X SSC 0.1% SDS, 0.2X SSC and 0.1X SSC and finally spin dried.

**Library preparation and sequencing**

Spatially barcoded ssDNA fragments were released from the surface by denaturation with 0.08N KOH at room temperature for 10 minutes and then quenched in 10 ul of 1M Tris pH7. The denatured fragments were pH adjusted with sodium acetate and cleaned with MinElute Reaction Cleanup Kit (Qiagen, 28204). The eluted DNA was then amplified using PCR using Partial.R1 and Ad2.short oligonucleotides for 15 cycles using PrimeSTAR Max DNA Polymerase mix (Takara, R045B). The amplified products were purified using 0.9X SPRI beads and i7-indexed in a second PCR (4 cycles) using PE1.0 and Ad2.X (where X is the sample index from [4]) oligonucleotides and KAPA HiFi HotStart Mix (Roche, KK2602). The final indexed libraries were cleaned up using 0.8X SPRI beads and adjusted to the desired molarity based on the concentrations measured using Qubit HS dsDNA Assay Kit (Thermo, Q32854) and the average fragment size from HS DNA Bioanalyzer kit (Agilent, 5067-4626).
Pooled libraries were then sequenced on Illumina Nextseq 550 or 2000 instrument using custom sequencing oligonucleotides for Read1 and Index2 (CustomR1 and CustomI2). We sequenced 65 bases for reads 1 and 2 (genomic sequence), 28 bases for i5 (spatial barcode and UMI), and 8 bases for i7 (sample index). All DNA oligonucleotides are listed in Table S1.

**H&E staining**

Tissue sections from breast cancer specimens were first dried with isopropanol (Fisher Scientific, A461-1) before staining. The sections were then stained with Mayer's hematoxylin (Agilent, S3309) for 4 minutes, washed in ultrapure water, incubated in bluing buffer (Agilent, CS702) for 2 minutes, washed in Milli-Q water, and further incubated for 1 minute in 1:20 eosin solution (Sigma-Aldrich, HT110216) in Tris-buffer (pH 6). The tissue sections were dried for 5 min at 37 °C and then mounted with 85% glycerol (Merck, 104094) and a coverslip. Imaging was performed using the Metafer VSlide system at ×20 magnification.

**Data pre-processing**

Raw reads were pre-processed using 10X Genomics' CellRanger ATAC pipeline (v2.0.0). We used a custom ′barcode_whitelist′ specifying positional barcodes from the spatial arrays and default reference genomes (mm10, v2.0.0 for the mouse data and hg38, v2.0.0 for the human data). All other parameters for ′mkfastq′ and ′count′ functions were set to default. Sequencing data from each section was processed separately and subsequently integrated with Seurat (v4.1.0, [5]) and Harmony (v0.1.0, [6]) R packages (*see below*).

**Analysis and visualisation**

For the embryos, we assayed sections across different developmental stages and integrated them for downstream analysis. To do so, we first obtained age-specific fragment files from ENCODE[7] and merged them using GenomicRanges's (v1.46.1,[8]) ´reduce()´ function. We then used these to create new accessibility matrices with a common set of 269,767 peaks. We also created new peak barcode matrices for the human breast cancer sample using a set of 215,978 peaks from [9]. We next subset the matrices to only include spots overlaying tissue, which were manually identified in Loupe Browser (v6.0.0) after aligning immunofluorescence pictures with capture area fiducials. Loupe browser was also used to select SOX2+ and SOX2- cortical spots in two E15.5 sections. The spatial object was created using STutility R package (v0.1.0,[10]), using tissue spot coordinates adjusted for the dimensions of the microscope images. STutility was used to produce the spatial plots using ´ST.FeaturePlot()´ function for quantitative variables. TSS enrichment plots and FragmentHistograms were generated using ArchR[11].

For each tissue type, we merged sections and performed normalization and dimensionality reduction on all peaks using Signac's (v1.6.0, [12]) ´RunTFIDF()´ and ´RunSVD()´ functions with default settings. We calculated gene activity using Ensembl annotations (EnsDb.Mmusculus.v79, v2.99.0 and EnsDb.Hsapiens.v86, v2.99.0), followed by log-normalization and principal component analysis (PCA). Genes from the Pcdh and Ugt gene clusters were removed from the gene activity assay prior to downstream analysis. For the embryos, graph clustering and UMAP were then performed on the peaks assay after Harmony integration on the top 7 dimensions and at a resolution of 0.7. Human breast cancer sections, which were obtained from the same tissue specimen, were merged directly using Seurat's ´merge()´ function followed by UMAP and graph clustering on dimensions 2 to 7, and at a resolution of 0.5. Cluster-wise Spearman's correlation of the chromatin accessibility profile was calculated for peaks around the transcription start site (i.e., between –1000 bp and +100 bp from TSS position) and for distal elements, using GenomicRanges's ´GetTSSPositions()´ followed by Signac's ´ClosestFeature()´ functions to annotate the peaks, and Seurat's ´AverageExpression()´ to obtain cluster-wise average accessibility levels for each peak. Differential accessibility analysis was performed on peaks using Seurat's ´FindAllMarkers()´ function with ´method = "LR"´ and unique fragments as the latent variable, and with ´logfc.threshold = 0.2´ and ´min.pct = 0.01´ to account for the sparsity of ATAC-seq data. ´FindAllMarkers()´ was also ran on the gene activity data with Wilcoxon Rank Sum test and followed by Gene Ontology analysis using gprofiler2 R package (v.0.2.1). Co-accessible peaks were identified by running the ´LinkPeaks()´ function on differentially accessible genes with a 0.05 pvalue and correlation cutoff as well as a minimum 1kb distance from the TSS. Motif enrichment analysis was performed by running ´FindMotifs()´ function using a set of clustered motifs from [13] on all linked peaks. Non-redundant top motifs were highlighted. For motif enrichment analyses in the developing mouse cortex, we first ran ´FoldChange()´ on peaks from SOX2+ and SOX2- cortical spots and then selected the top 500 peaks for motif analyses as above. Full lists of enriched motifs are provided in Table S2. Vista enhancers were downloaded from https://enhancer.lbl.gov/ and genome coordinates were lifted to mm10 using UCSC liftover tool prior to intersection with spatial ATAC tissue-specific peaks using bedtools [14].

**Denoising**

Using a deep count autoencoder, DCA (v0.3.4,[15]), we denoised the peak barcode matrix of the combined object, as well as the gene activity matrices. For the peaks data, we specified the following parameters: ´--nosizefactors --nonorminput --nologinput´, whereas DCA was run with default settings on the gene activity data. Additionally, we performed DCA with default parameters on Visium data from the mouse embryo and human breast cancer (see below). Dimensionality reduction and clustering was performed on the denoised data as above to evaluate concordance between original and denoised data. While clustering and differential accessibility analysis was performed on original data, denoised data were used for visualisation of accessibility levels and for multimodal integration with single cell data (*see below*).

**Spatial analysis**

STutility's ´RunNMF()´ function was run with ´nfactors =8´ after ordering the top 25% variable features according to spatial correlation. Harmony integration and graph clustering was performed using NMF's factors in dimensionality reduction and the groups obtained this way were compared with the spatial-agnostic clustering results obtained from the original peaks assay.

**Visium**

The 10X Genomics' Visium platform was used to obtain spatial transcriptomics data for tissue samples matching our spatial ATAC sections (e.g., either on consecutive tissue slices from breast tumor block or on similar sagittal level of embryos from the same litter).
Raw data were pre-processed using SpaceRanger's (v1.3.1) ´mkfastq´ and ´count´ functions with default parameters and the resulting gene-barcode matrices were then analysed with Seurat for normalization, dimensionality reduction and clustering, and with STUtility for plotting. Visium data were denoised with DCA and default parameters for visualizations and comparison with spatial ATAC data.

**Integrative multimodal analysis**

We performed multimodal comparison of our spatial ATAC data using either spatial or single-cell transcriptomics. To measure cluster-wise concordance between gene expression and accessibility, we analysed in parallel spatial ATAC and spatial RNA-seq data from the embryos and obtained cluster markers for each modality, which we used to calculate module scores (with Seurat's ´AddModuleScore()´) in each assay.
Additionally, we performed multimodal integrative analysis between spatial ATAC and single-cell RNA-seq data. For the embryos, we restricted our analysis to the cortex of E15 mice and manually subset spots overlaying the region of interest. In parallel, we obtained a developmental transcriptional atlas from [16], and subset it to include cells from E15 brains. Furthermore, we restricted our analysis to only comprise the dorsal forebrain and specifically looked at cells in the neurogenic trajectory (i.e., radial glia, intermediate progenitors, and neurons). Single-cell data were processed according to Seurat's standard workflow and subset to n=1500 cells randomly sampled across the clusters. We integrated spatial ATAC and single-cell RNA-seq data using canonical correlation analysis (CCA) and 2000 anchor features. Co-embedded data were subsequently subjected to dimensionality reduction using PCA. UMAP visualizations calculated on the top 7 components were, finally, used to order cells in pseudotime using monocle3 (v1.0.0,[17]) and the radial glia cluster as root cells.
For the human breast cancer data, we obtained a comprehensive single-cell RNA-seq atlas [18] and processed it with Seurat's standard workflow. We then probed enrichment of the major cell types in our spatial ATAC and spatial RNA-seq clusters. To do so, we adopted the author's classification of cells in the highest tier (i.e., "celltype_major") and used Seurat's label transfer workflow based on CCA to obtain prediction scores for each cell type in the single cell dataset.

**Code and data availability**

All raw data and processed count matrices can be accessed at GSEXXX. All analysis code used can be found at https://github.com/marzamKI/spatial_atac

**Table S1.** Oligonucleotides sequences

**Table S2.** Differentially accessible gene and peak sets. Motif enrichment analyses. Public datasets used.

**Table S3.** Patient data and tumor characteristics

### Methods references


1. Schindelin, J. *et al.* Fiji: an open-source platform for biological-image analysis. *Nat Methods* **9**, 676–682 (2012).
2. Picelli, S. *et al.* Tn5 transposase and tagmentation procedures for massively scaled sequencing projects. *Genome Res.* **24**, 2033–2040 (2014).
3. Corces, M. R. *et al.* An improved ATAC-seq protocol reduces background and enables interrogation of frozen tissues. *Nature methods* **14**, 959–962 (2017).
4. Buenrostro, J. D., Giresi, P. G., Zaba, L. C., Chang, H. Y. & Greenleaf, W. J. Transposition of native chromatin for fast and sensitive epigenomic profiling of open chromatin, DNA-binding proteins and nucleosome position. *Nature methods* **10**, 1213–1218 (2013).
5. Hao, Y. *et al.* Integrated analysis of multimodal single-cell data. *Cell* **184**, 3573-3587.e29 (2021).
6. Korsunsky, I. *et al.* Fast, sensitive and accurate integration of single-cell data with Harmony. *Nat Methods* **16**, 1289–1296 (2019).
7. Gorkin, D. U. *et al.* An atlas of dynamic chromatin landscapes in mouse fetal development. *Nature* **583**, 744–751 (2020).
8. Lawrence, M. *et al.* Software for Computing and Annotating Genomic Ranges. *PLOS Computational Biology* **9**, e1003118 (2013).
9. Corces, M. R. *et al.* The chromatin accessibility landscape of primary human cancers. *Science* **362**, eaav1898 (2018).
10. Bergenstråhle, J., Larsson, L. & Lundeberg, J. Seamless integration of image and molecular analysis for spatial transcriptomics workflows. *BMC Genomics* **21**, 482 (2020).
11. Granja, J. M. *et al.* ArchR is a scalable software package for integrative single-cell chromatin accessibility analysis. *Nat Genet* **53**, 403–411 (2021).
12. Stuart, T., Srivastava, A., Madad, S., Lareau, C. A. & Satija, R. Single-cell chromatin state analysis with Signac. *Nat Methods* **18**, 1333–1341 (2021).
13. Vierstra, J. *et al.* Global reference mapping of human transcription factor footprints. *Nature* **583**, 729–736 (2020).
14. Quinlan, A. R. & Hall, I. M. BEDTools: a flexible suite of utilities for comparing genomic features. *Bioinformatics* **26**, 841–842 (2010).
15. Eraslan, G., Simon, L. M., Mircea, M., Mueller, N. S. & Theis, F. J. Single-cell RNA-seq denoising using a deep count autoencoder. *Nat Commun* **10**, 390 (2019).
16. La Manno, G. *et al.* Molecular architecture of the developing mouse brain. *Nature* **596**, 92–96 (2021).
17. Cao, J. *et al.* The single-cell transcriptional landscape of mammalian organogenesis. *Nature* **566**, 496–502 (2019).

18. Wu, S. Z. *et al.* A single-cell and spatially resolved atlas of human breast cancers. *Nat Genet* **53**, 1334–1347 (2021).
